# Defining the adult hippocampal neural stem cell secretome: in vivo versus in vitro transcriptomic differences and their correlation to secreted protein levels

**DOI:** 10.1101/760215

**Authors:** JK. Denninger, X. Chen, AM. Turkoglu, P. Sarchet, AR. Volk, P. Yan, ED. Kirby

## Abstract

Recent evidence shows that adult hippocampal neural stem and progenitor cells (NSPCs) secrete a variety of proteins that affect tissue function. Though several individual NSPC-derived proteins have been shown to impact cellular processes like neuronal maturation and stem cell maintenance, a broad characterization of NSPC-secreted factors is lacking. Secretome profiling of low abundance stem cell populations is typically achieved via proteomic characterization of in vitro, isolated cells. Here, we analyzed the in vitro NSPC secretome using conditioned media from cultured adult mouse hippocampal NSPCs and detected over 200 different bioactive proteins with an antibody array. We next assessed the NSPC secretome on a transcriptional level with RNA sequencing (RNAseq) of cultured NSPCs. This comparison revealed that quantification of gene expression did not accurately predict relative protein abundance for several factors. Furthermore, comparing our transcriptional data with previously published single cell RNA sequencing datasets of freshly isolated hippocampal NSPCs, we found key differences in gene expression of secreted proteins between cultured and acutely isolated NSPCs. Understanding the components and functions of the NSPC secretome is essential to understanding how these cells may modulate the hippocampal neurogenic niche, as well as how they can be applied therapeutically. Cumulatively, our data emphasize the importance of using proteomic analysis in conjunction with transcriptomic studies and highlights the need for better methods of global unbiased secretome profiling.

## 1. Introduction

In the adult mammalian brain, neural stem cells (NSCs) are located in two specific niches, the subventricular zone (SVZ) and the subgranular zone (SGZ) of the hippocampal dentate gyrus (DG) (Gage and Temple 2013). In the adult SGZ, resident NSCs produce neuronal intermediate progenitor cells (nIPCs) that proliferate throughout the lifespan of the organism to produce functionally-relevant neurons and astrocytes (Ming and Song 2011). In addition to creating mature cell types, NSCs also secrete an array of growth factors and cytokines, collectively termed the stem cell secretome (Drago et al. 2013). In the healthy adult SGZ, NSCs and their progenitors (together NSPCs) have been shown to produce the secreted factors milk-fat globule EGF-factor 8 (MFGE8) and vascular endothelial growth factor (VEGF), both of which regulate NSPC maintenance through autocrine signaling (Kirby et al. 2015; Zhou et al. 2018), as well as PTN which drives maturation of immature, developing neurons (Tang et al. 2019). However, while the secretome of other tissue stem cells, such as mesenchymal stem cells, has been extensively catalogued (Liang et al. 2014), a comprehensive characterization of the NSPC secretome is lacking (Andres et al. 2011; Ryu et al. 2004; Yasuhara et al. 2006; Ourednik et al. 2002; Tang et al. 2019).

Protein-level analysis of the stem cell secretome is typically accomplished using in vitro models, where the conditioned media (CM) of isolated stem cells is profiled using proteomics approaches such as antibody-based arrays. While this approach has provided the first broad understanding of the content and variety of several tissue stem cell secretomes (Skalnikova et al. 2011), it is limited in scope to the pre-selected factors present on the arrays. Furthermore, comparison of cultured cells to their in vivo counterparts in several organ systems has provided increasing evidence of significant changes that occur when cells are transitioned from an in vivo to an in vitro environment (Durr et al. 2004; Binato et al. 2013; Duggal et al. 2009). These changes were not only dependent on the method of isolation (Wernly et al. 2017), but also the conditions of culture maintenance (Shahdadfar et al. 2005; Duggal and Brinchmann 2011). More specific to NSPCs, key differences in gene expression of inflammatory and cytokine signaling factors were identified in cultured SVZ NSCs versus NSCs acutely isolated from the SVZ (Dulken et al. 2017).

High-throughput characterization of tissue stem cell secretomes in vivo faces several challenges. In vivo, adult tissue stem cells, including hippocampal NSCs, are small populations residing in heterogeneous niches that impede sufficient isolation of cells that are necessary for most large-scale proteomics protocols. Methods such as immunostaining of tissue sections or flow cytometry of dissociated cells allow detection of pre-selected proteins in specific cells, but are limited in the total number and type of proteins that can be examined in each instance. To broaden the scope of in vivo protein investigation, extracellular factors can be identified from tissue niches with high performance liquid chromatography (HPLC) or mass spectrometry of microdialysates. However, these methods fail to determine the cells that produce the secreted factors.

In contrast to proteomics approaches, high throughput RNA sequencing (RNAseq) is adaptable to low cell input and several studies have recently provided large-scale quantification of gene expression in acutely isolated NSPCs by single cell RNAseq (scRNAseq) (Hochgerner et al. 2018; Shin et al. 2015; Artegiani et al. 2017; Dulken et al. 2017; Zywitza et al. 2018). The emergence of scRNAseq provides a method for assessing the putative adult DG NSPC secretome at the mRNA expression-level without the perturbations inherent in culturing cells in vitro. However, transcriptional activity does not always correlate well with bioavailable protein due to added layers of regulation on translational and post-translational levels (Lipshitz, Claycomb, and Smibert 2017; Besse and Ephrussi 2008).

Given that mRNA levels may not reflect secreted protein levels and in vitro gene expression may not mirror in vivo expression, we chose to create a multi-faceted analysis of the adult DG NSPC secretome from mRNA level to secreted protein in both in vitro and in vivo models. For high-throughput protein-level quantification, we used a protein array to probe for over 300 different cytokines and growth factors secreted by cultured adult hippocampal NSPCs. Then, to assess the correlation between mRNA and secreted protein, we determined the relative transcriptional expression of the highest secreted proteins using RNA sequencing of cultured NSPCs. To evaluate the equivalence of cultured and endogenous NSPCs, we examined two independently published single cell sequencing datasets of hippocampal NSPCs (Shin et al. 2015; Hochgerner et al. 2018). Comparison of transcript expression for secreted factors identified in our protein array not only demonstrated important divergence between transcriptional and translational expression, but also exposed significant differences between cultured and in vivo hippocampal NSPCs. Our findings therefore not only provide a broad characterization of the putative adult DG NSPC secretome, but also highlight the need to consider limitations of currently available methods for secretome profiling.

## 2. Results

### 2.1. Protein array of cultured NSPC conditioned media detected several secreted factors that impact NSPC function

Conditioned media from three separate adult hippocampal NSPC cultures (one male and two female) were collected and probed for over 300 growth factors, cytokines, chemokines, adipokines, angiogenic factors, proteases, soluble receptors and soluble adhesion molecules with a protein antibody array. Supplemental Table 1 lists the fluorescence values after normalization to average fluorescence intensity within each slide for all factors that were detected in the assay ranked by Z-score. Of these, the top 50 most highly detected factors were selected for further scrutiny (Fig.1A). These factors included VEGF, which we previously identified as highly secreted by NSPCs and critical for stem cell maintenance in vivo and in vitro (Kirby et al. 2015). In addition, many other soluble growth factors and cytokines were detected, along with several carrier proteins that are implicated in NSPC maintenance such as those of the IGFBP family (IGFBP3, IGFBP2, and IGFBP5) (Kalluri and Dempsey 2011; Shen et al. 2019; Barkho et al. 2006). Soluble fragments of receptors that promote NSPC maintenance and proliferation, such as TYRO3, were also found, likely reflecting shedding of the extracellular portions from these typically membrane-integral receptors (Ji et al. 2014). Known regulators of apoptosis that promote NSPC survival, FAS and SPP1, were also present in relatively high quantities (Knight, Scharf, and Mao-Draayer 2010; Rabenstein et al. 2015). Interestingly, CD70 on NSPCs has been shown to interact with CD27 on CD4+ T-cells to induce FAS-mediated apoptosis of T cells, but in our array, we detected CD27 as being secreted by NSPCs which may imply an additional layer of control in NSPC modulation of immune cells (Lee et al. 2013). Cumulatively, the protein array confirms the presence of some previously reported components of the NSPC secretome and suggests numerous other potentially bioactive proteins produced and secreted by these cells.

**Fig. 1.**
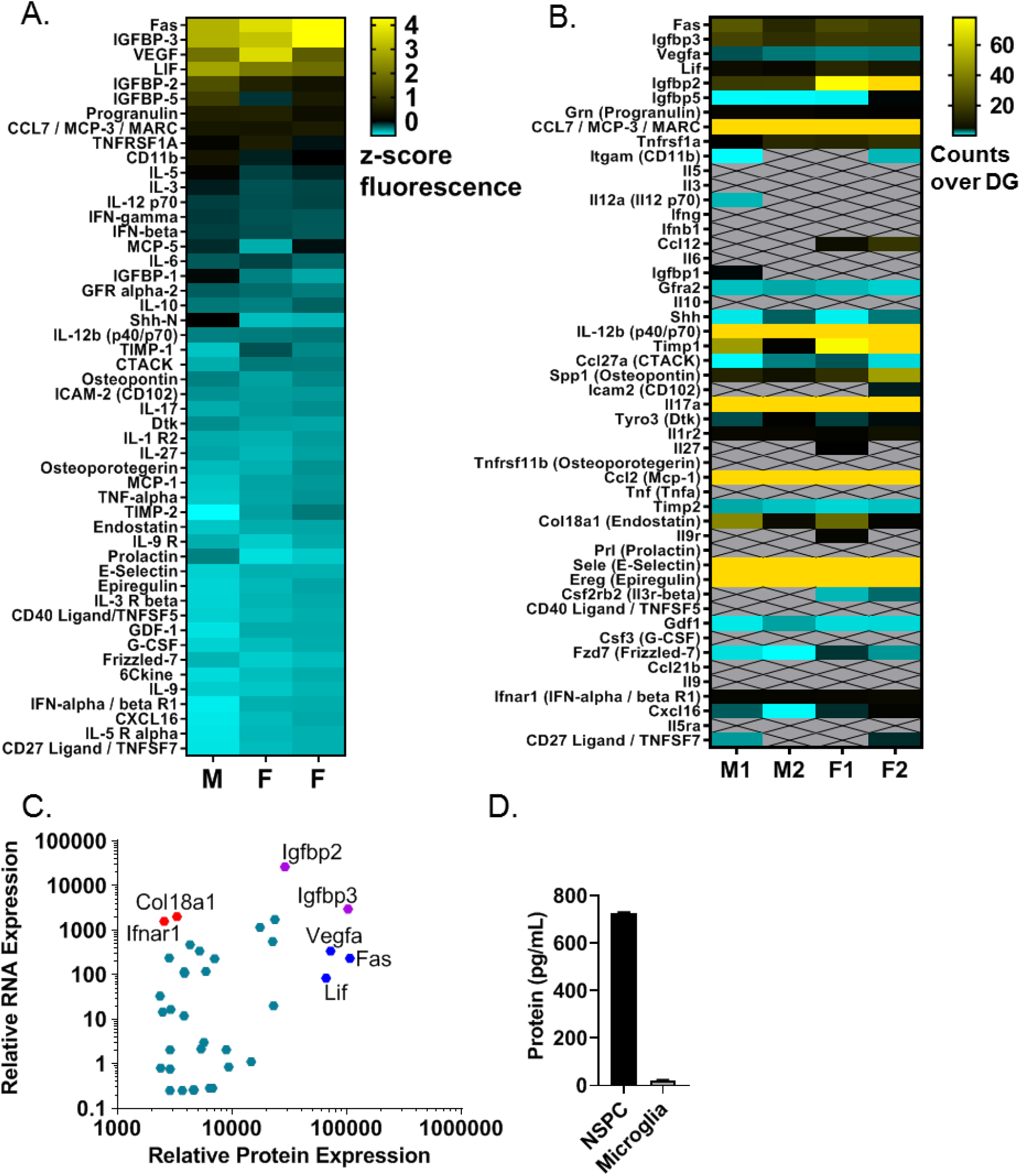
Top 50 secreted factor analysis in cultured NSPCs. (A) Heatmap of top 50 proteins detected in 48h conditioned NSPC media. Data normalized to average fluorescence intensity within each slide before Z-score transformation. (M = male, F = female). (B) Heatmap of transcript expression from RNAseq of cultured NSPCs for the 50 proteins identified in protein array. Data normalized to whole DG. (M = male, F = female). (C) Scatter plot of relative transcript expression compared to relative protein expression for the top 50 proteins identified in protein array. Protein expression is represented as the fluorescence intensity normalized to the average fluorescence intensity within the slide. RNA expression represented as total counts. (D) VEGF protein concentration in NSPC conditioned media after 48h of culture measured by VEGF ELISA. Conditioned media from microglia is provided as a comparison control population which is expected to secrete low levels of VEGF (n = 4).

### 2.2. RNA sequencing of cultured NSPCs revealed that transcriptional expression does not strongly equate to relative protein expression of secreted factors

Four separate cultures of adult hippocampal NSPCs were harvested and analyzed by RNA sequencing. A total of 50600 unique transcripts were detected (Supplemental Table 2). Total counts in RNAseq of the cultured NSPCs for the genes that encode the top 50 secreted factors identified in the above protein array were normalized to the total counts for those genes in RNAseq of the whole DG (Fig. 1B). Comparing mRNA expression in cultured NSPCs relative to whole DG first revealed that the majority of the top 50 protein factors from conditioned media were not enriched compared to whole DG at the mRNA level. This suggests either that transcriptional activity does not predict final protein secretion or that other DG cell types also express large quantities of these transcripts such that NSPCs do not show enrichment compared to whole tissue expression. Almost half of the factors were not detected in any of the samples, implying either transcript expression that is below the threshold for detection by RNAseq or false positive detection in the protein array. Either possibility poses problems for using one method alone to identify the biologically-meaningful NSPC secretome and necessitates further assessment of those individual factors by additional methods.

To better determine whether mRNA predicts relative protein abundance independent of DG enrichment status, Spearman correlation analysis of relative protein levels versus relative transcript levels from cultured NSPCs for all 50 factors was performed (Fig. 1C). This analysis showed a moderate degree of correlation overall (r(48) = 0.3299, p = 0.0193). Though some factors, such as *Igfbp3* and *Igfbp2* (displayed in purple), displayed relatively high levels of both transcript and secreted protein detected in the conditioned media array, several factors, such as *Fas, Vegfa*, and *Lif* (displayed in blue) or *Col18a1* and *Ifnar1* (displayed in red) respectively, showed unexpectedly low or high transcriptional expression for their ranking at the secreted protein-level. For example, mRNA levels for *Vegfa* were not particularly high in RNAseq of cultured NSPCs, even though VEGF was the third highest secreted factor identified in the protein array. To confirm the array findings with a secondary method, we used a well-validated ELISA and found that cultured NSPCs secreted large amounts of VEGF, as much as 700 pg/mL (Fig. 1D). *Timp1*, in contrast, showed high mRNA expression, second only to *Igfbp2*, despite being 23^rd^ in the ranking of proteins. Thus while mRNA suggests TIMP1 is a prominent secreted product of NSPCs in vitro, the secreted protein quantification suggests more modest production of this factor. Notably, TIMP1 is a sex-specific factor and both the protein array and RNAseq show elevated levels of TIMP1 in the female samples compared to the male samples (Fig. 1A and B), suggesting that both methods are accurately detecting TIMP1.

### 2.3. Comparison of transcript levels for top 50 highest secreted proteins in cultured versus acutely isolated NSPCs indicated moderate overall correlation, but important divergence in individual genes

To explore how in vitro culturing might impact the NSPC secretome, we next compared our in vitro NSPC RNAseq with two previously published scRNAseq studies of adult hippocampal NSPCs (Shin et al. 2015; Hochgerner et al. 2018). For the Hochgerner et al. dataset (Hoch.), single cell data for radial glial like cells (RGLs, which correspond to NSCs), nIPCs, and neuroblasts (NBs, representing late neuronal progenitor cells), were averaged to generate an overall transcript count for each cell population. For the Shin et al. dataset, which identified single cells as representing points in pseudotime from a quiescent stem cell to an activated intermediate progenitor, NSC and nIPC groups were created such that NSCs were represented by the averaged counts of all cells between pseudotime points 0 and 0.479, while nIPCs were averaged counts for pseudotime points between 0.506 and 1. Comparison of mRNA for the top 50 secreted proteins in vitro with these in vivo NSPC subpopulations revealed no correlation across datasets (Fig. 2A and B) as well as failure to detect several genes on an RNA level (Fig. 3).

**Fig. 2.**
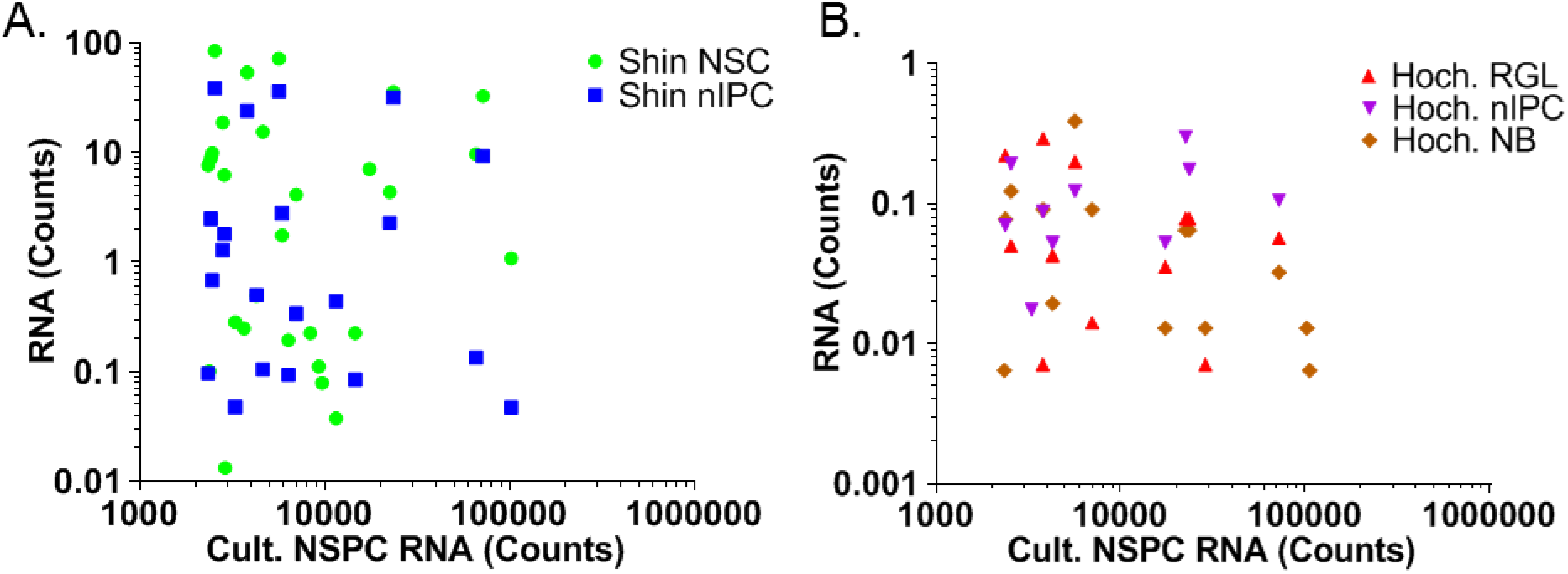
Comparison of in vivo versus in vitro transcript level expression of top 50 secreted factors identified in protein array. (A) Scatter plot of relative transcript expression in Shin NSCs and nIPCs compared to cultured (Cult.) NSPCs. (B) Scatter plot of relative transcript expression in Hochgerner (Hoch.) RGLs, nIPCs, and NBs compared to cultured NSPCs. All RNA expression represented as averaged counts for all cells within the population.

**Fig. 3.**
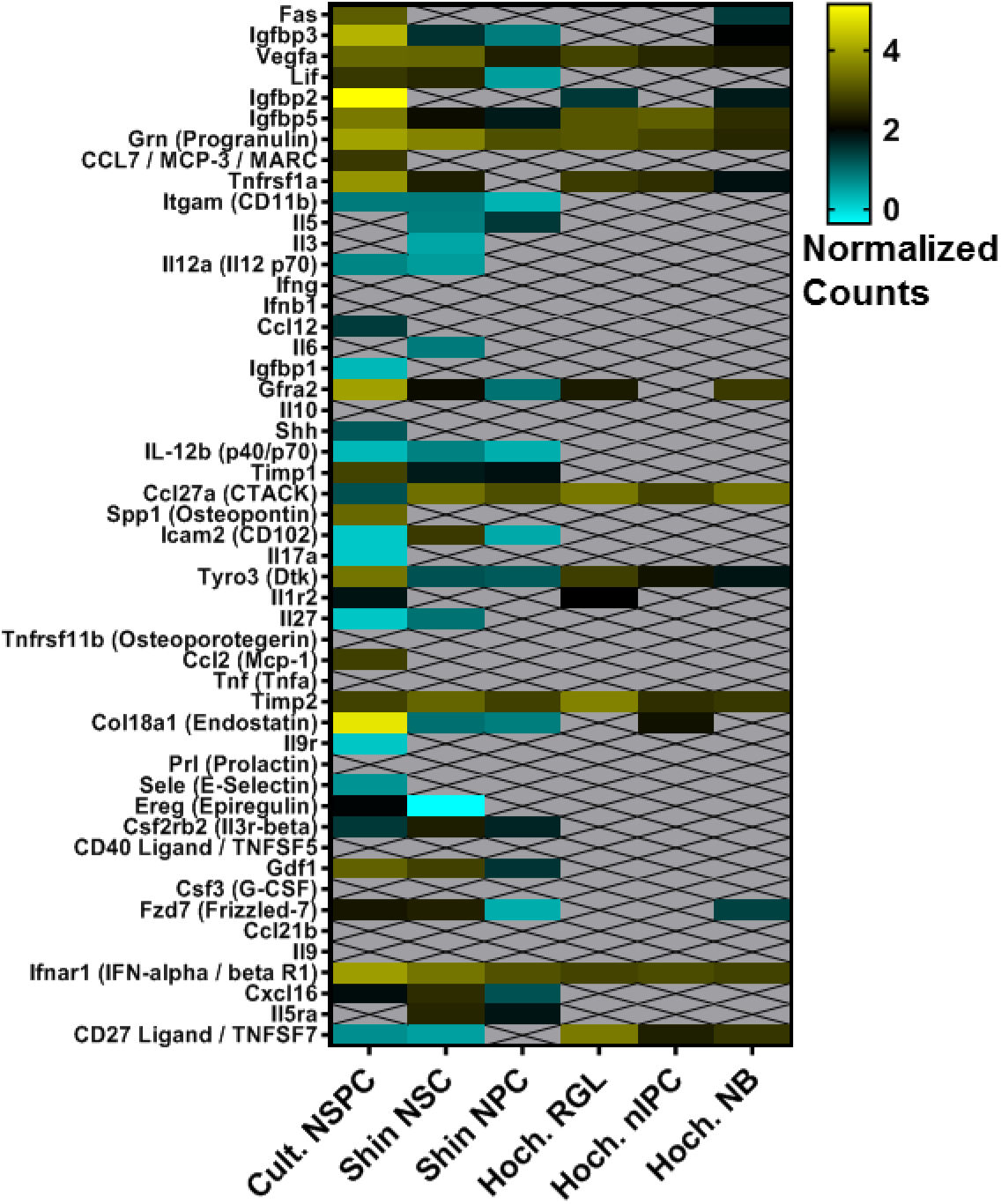
Heatmap of in vivo and in vitro transcript level expression of top 50 secreted factors identified in protein array. All RNA expression represented as the log_10_ transformed value of *Actb* normalized counts over total counts per sample x 10^3^.

First, comparison of log transformed *Actb* normalized counts across all datasets showed that several genes were not detected in either of the scRNAseq datasets, likely due to limitations in input RNA, total cell number, and sequencing depth. Many of the lowly transcribed genes in the cultured (Cult.) NSPCs were only detected in Shin scRNAseq dataset, which mirrored the relatively low in vitro expression (Fig. 3). Several of the relatively highly transcribed genes in vitro, such as *Grn, Igfbp5*, and *Ifnar1*, that had similar relatively high levels of transcription in vivo. Other genes, such as *CD27, Gfra2, Ccl27a*, and *Col18a1* showed opposing relative transcript levels (Fig. 3).

As the division of in vivo RNAseq data in to subpopulations implies, NSPCs are heterogeneous. NSPCs exist mostly as NBs with a substantial minority existing as RGLs/NSCs (Encinas and Enikolopov 2008). The nIPC state represents a relatively short-lived transition and therefore small population overall (Hochgerner et al. 2018; Encinas and Enikolopov 2008). Given these relative population distributions, Spearman correlations between cultured NSPCs and in vivo subpopulations were expected to vary. Specifically, we expected correlations between in vitro and in vivo data to be strongest with NBs from Hochgerner, followed by Hochgerner RGLs, mirroring the expected size of these subpopulations in NSPCs as a whole. The weakest relationships were expected between cultured NSPCs, Shin nIPCs and Hochgerner nIPCs because nIPCs are a small contributor to the NSPC population. Consistent with this prediction, within the Shin dataset, we found a stronger, but still not statistically significant, correlation between the cultured NSPCs and Shin NSCs (r(48) = 0.05176, p = 0.7211) than with Shin nIPCs (r(48) = 0.04415, p = 0.7608) (Fig. 2A). Hochgerner et al. data also showed stronger correlation of cultured NSPCs with NBs (r(48) = 0.2134, p = 0.1368) than with nIPCs (r(48) = 0.08846, p = 0.5413). The correlation with NBs was also greater than with RGLs (r(48) = 0.1433, p = 0.3209) (Fig. 2B), again consistent with the expected relative populations in cultured NSPCs.

### 2.4. Global comparison of cultured versus acutely isolated NSPC transcriptomes illustrated differences in gene expression in vitro versus in vivo

To expand the analysis of in vitro versus in vivo secretome beyond the proteins detected by the antibody array, we next performed Gene Ontology (GO) analysis of transcripts detected above zero in in vitro NSPCs to identify 313 genes that encode proteins located in the extracellular region (Supplemental Table 3). Relative transcript expression normalized to the whole DG for these factors was consistent in each of the four cultured NSPC samples (Fig. 4). Excluding genes that failed to generate transcripts, comparison of transcript levels in cultured NSPCs to the Shin dataset showed moderate correlation with NSCs (r(265) = 0.3710, p < 0.0001) and nIPCs (r(234) = 0.2419, p < 0.0001) (Fig. 5A). Similarly, comparison with the Hochgerner dataset showed moderate correlation with RGLs (r(191) = 0.4490, p < 0.0001), nIPCs (r(170) = 0.3301, p < 0.0001) and NBs (r(187) = 0.2824, p < 0.0001) (Fig. 5B).

**Fig. 4.**
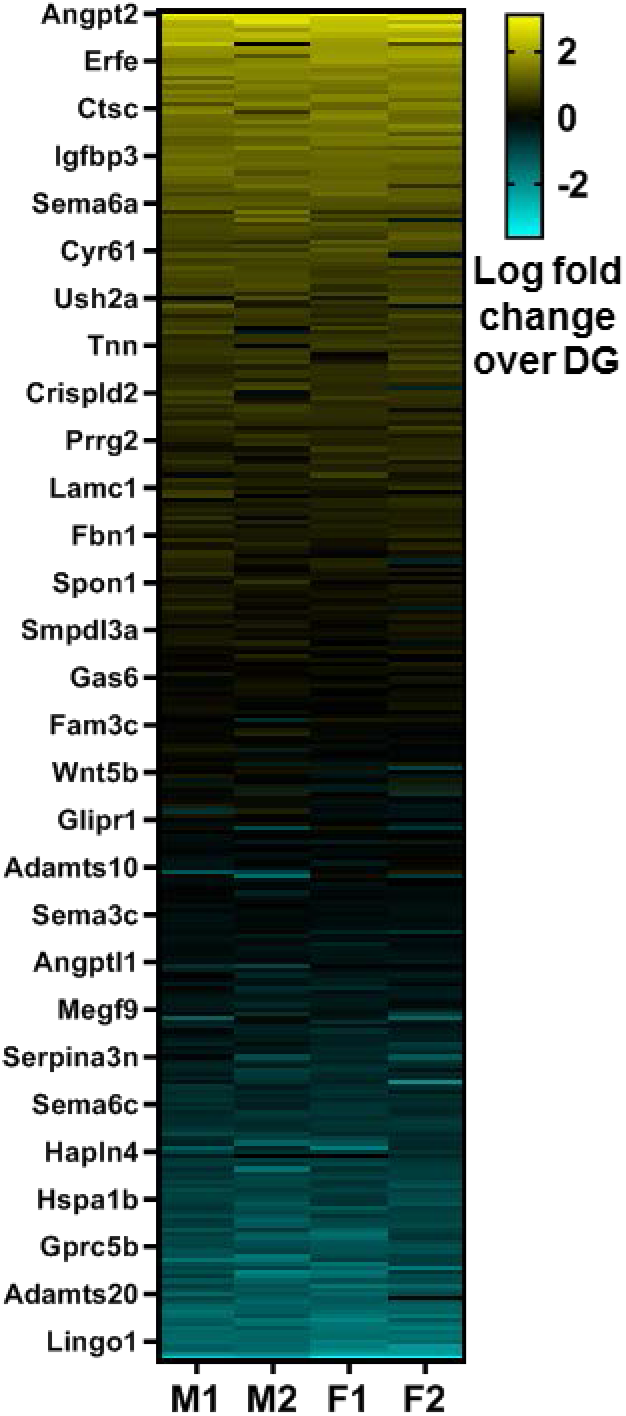
Heatmap of transcript expression for extracellular proteins identified by GO analysis of all transcripts above zero in RNAseq of cultured NSPCs. Data represented as the log_10_ transformed counts normalized to whole DG. (M = male, F = female).

**Fig. 5.**
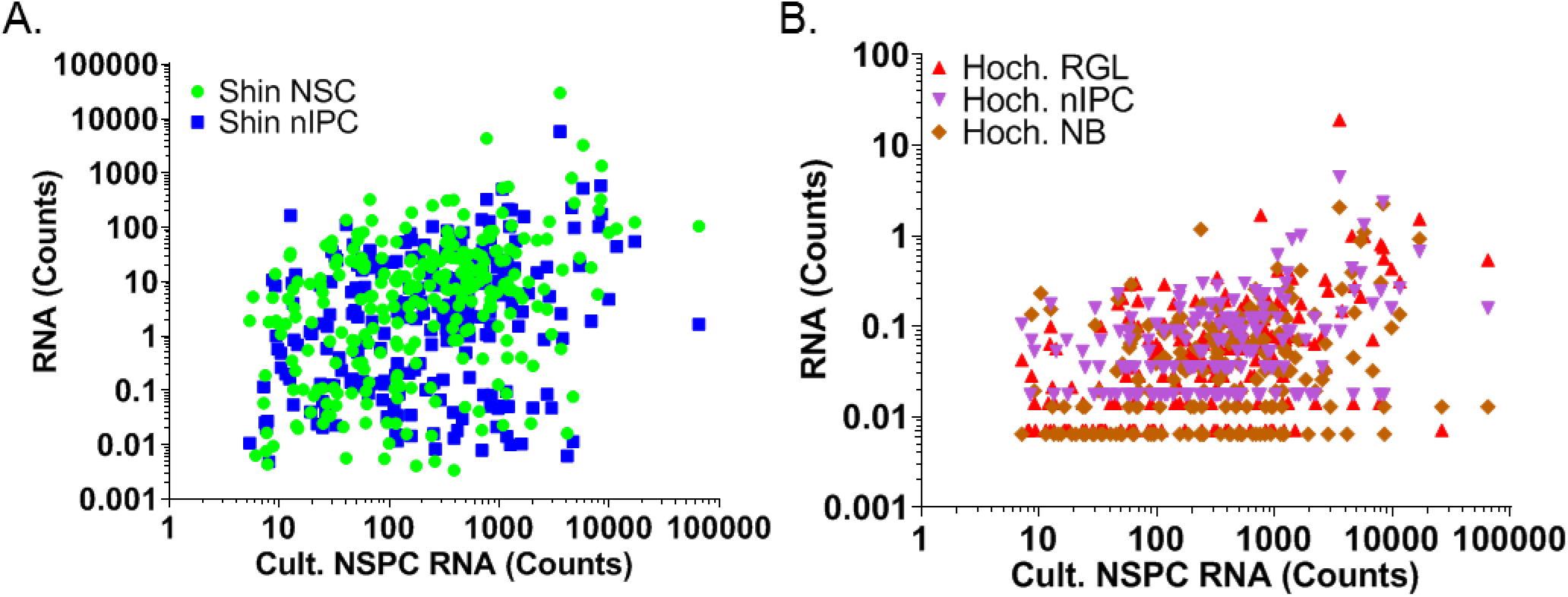
Comparison of in vivo versus in vitro transcript level expression for extracellular proteins identified in RNAseq of cultured NSPCs. (A) Scatter plot of relative transcript expression in Shin NSCs and nIPCs compared to cultured NSPCs. (B) Scatter plot of relative transcript expression in Hochgerner RGLs, nIPCs, and NBs compared to cultured NSPCs. All RNA expression represented as averaged counts for all cells within the population.

We next expanded the comparison of transcriptional gene expression in vitro versus in vivo beyond secreted proteins to gain a more unbiased appreciation of how culturing may alter NSPCs. Comparison of the top 1000 cultured NSPC transcripts with the expression found in the Shin dataset indicated low correlation with NSCs (r(938) = 0.1919, p < 0.0001) and nIPCs (r(931) = 0.1727, p < 0.0001) (Fig. 6A). Likewise, comparison with the Hochgerner dataset revealed low correlation between in vitro NSPCs and in vivo RGLs (r(917) = 0.09464, p = 0.0041), nIPCs (r(913) = 0.1441, p < 0.0001), and NBs (r(915) = 0.1448, p < 0.0001) for these genes (Fig. 6B). Thus, comparison of genes that are not limited to extracellular proteins confirm an increased divergence in transcription between in vitro and in vivo cells.

**Fig. 6.**
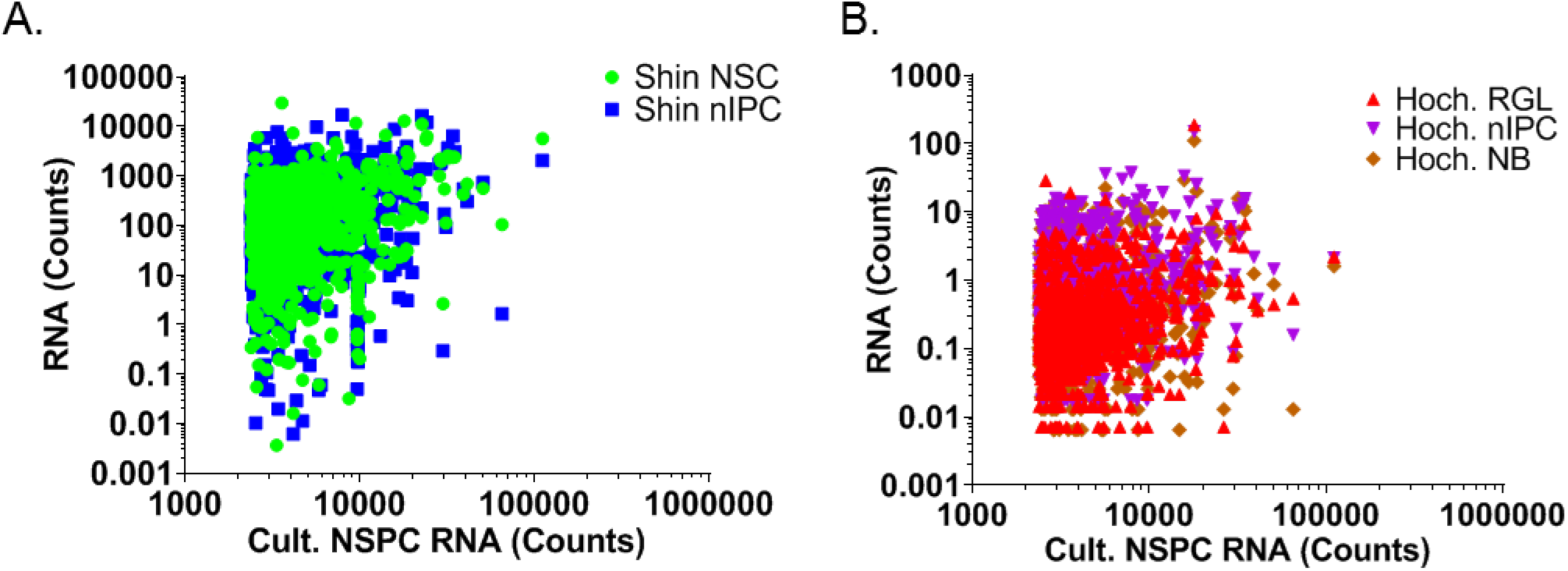
Comparison of in vivo versus in vitro transcript level expression for top 1000 most highly transcribed genes in cultured NSPCs. (A) Scatter plot of relative transcript expression in Shin NSCs and nIPCs compared to cultured NSPCs. (B) Scatter plot of relative transcript expression in Hochgerner RGLs, nIPCs, and NBs compared to cultured NSPCs. All RNA expression represented as averaged counts for all cells within the population.

## 3. Discussion

Over the past decade, stem cell secretomes have been investigated for their potential therapeutic applications to various neurologic disorders (Drago et al. 2013; Baraniak and McDevitt 2010; Liang et al. 2014). The cytokines, chemokines, and growth factors that comprise the neural stem cell secretome have been thus far identified individually or in limited combinations with traditional methods of protein analysis such as immunohistochemistry, western blotting, and ELISA (Skalnikova et al. 2011). In other tissue stem cells, broader characterization of secreted proteins has been achieved with array based methods examining isolated stem cell populations maintained in vitro (Skalnikova et al. 2011). More recently, in vivo stem cell profiling with low cell and scRNAseq has emerged as a proxy for untargeted protein level analysis in small populations like NPSCs that do not yield sufficient protein for high-throughput translational-level analysis (Artegiani et al. 2017; Hochgerner et al. 2018; Shin et al. 2015). Cumulatively, these methods have provided several informative pieces of the puzzle. Our findings at the transcriptional level in vitro and in vivo, as well as the protein-level in vitro, provide a first attempt at a comprehensive analysis of the NSC secretome, and, in the process, highlight significant limitations to current approaches.

First, we found that transcriptional-level analysis via RNAseq does not serve as a reliable surrogate for global, unbiased protein expression analysis. Using an array of over 300 preselected proteins, we showed that for the top 50 secreted factors detected in cultured NSPCs, mRNA transcription only moderately correlated to protein expression. Proteins known to significantly impact NSPC function and derive in measurable quantities from NPSCs in vivo (i.e. VEGF) (Kirby et al. 2015) showed considerable divergence in RNA versus protein levels, implying that had RNAseq been the sole initial screening method to identify important NSPC factors, VEGF would not have been included. Thus, these data elucidate important discrepancies between RNA and protein expression and also highlight the importance of a multifaceted approach involving protein-level and transcript-level analyses to identify secretome components. As part of this multifaceted approach to NSPC characterization, a method to identify the NPSC proteome in an unbiased manner is needed, but currently lacking. Notably, though, advances in using noncanonical amino acids and transgenic tRNA synthetases to label proteins cell-specifically have recently been adapted to mammalian systems to execute such global proteomic analysis (Mahdavi et al. 2016; Liu et al. 2017; Elliott, Bianco, and Chin 2014). While this approach has been successfully applied to in vivo mature neuronal populations (Alvarez-Castelao et al. 2019), it has yet to be applied to adult stem cell populations.

Second, our data show significant changes in DG NSPC gene expression secondary to culturing, as has been reported for other stem cell populations (Binato et al. 2013; Duggal et al. 2009; Duggal and Brinchmann 2011; Wernly et al. 2017; Shahdadfar et al. 2005). For example, NSCs isolated from the SVZ displayed important differences in transcription of inflammatory cytokines compared to their in vivo counterparts (Dulken et al. 2017). Similarly, we found no significant agreement in RNA transcript levels for the top 50 in vitro secreted proteins between cultured DG NSPCs and hippocampal NSCs, nIPCs or NBs from previously published scRNAseq datasets (Shin et al. 2015; Hochgerner et al. 2018). The concordance between in vitro and in vivo transcript levels improved only modestly as more genes were included in the analysis, indicating significant divergence in the transcriptomes of endogenous versus cultured cells.

Though the disconnect between in vitro and in vivo gene expression may at first suggest that only in vivo studies should be pursued, such a focused approach does not faithfully represent all biologically and therapeutically relevant stem cell sources. Current clinical trials use cultured neural stem cell lines while preclinical models use NSPCs with varying amounts of in vitro processing (Mazzini et al. 2019; Ottoboni, Merlini, and Martino 2017; Mendes-Pinheiro et al. 2018; Chandanala et al. 2014). Thus, analysis of both cultured and acutely isolated stem cells is critical for the development of effective NSPC-based therapies. An open question here is whether the NSPC secretome after transplant in clinical/preclinical models resembles that of cultured cells or of in vivo cells.

Our in vitro RNAseq versus secreted protein analysis suggests that transcriptional expression and protein secretion is not equivalent. However, each method used in our studies comes with potential technical artefacts. Antibody arrays are a standard screening technique for stem cell secretomes (Skalnikova et al. 2011), but they may yield both false positive and false negative signals. Ideally, factors identified in these screens should undergo additional rigorous validation by targeted methods of cell-specific protein analysis. For our array, we verified one key protein, VEGF, with a well-established ELISA approach and found that this high-ranked protein in the array was also present in substantial absolute quantities by ELISA. We also noted that TIMP1, which is a sex-linked gene expressed more prominently in females, was consistently higher in samples derived from females than from males. Still, it is possible that some individual proteins from an array of this size were incorrectly quantified. In contrast, RNAseq is not subject to antibody specificity challenges. Instead, particularly in the case of scRNAseq, it can suffer from transcript drop-outs and RNA degradation, which can impair the robustness of the final data (Bhargava et al. 2015; Brennecke et al. 2013; L. Lun, Bach, and Marioni 2016). Furthermore, while our transcriptional data was normalized to transcript expression in the whole DG to account for endogenous gene expression by other cell types, the protein array has no analogous comparison that takes into account protein production by other cell types in the DG. Thus, proteins that were secreted in relatively high amounts by NSPCs may not be biologically relevant in vivo when compared to overall levels in the DG. These sources of imprecision are a hazard of high throughput approaches and underscore the need to further validate any one target protein if it is to be pursued for therapeutic or investigational purposes.

In conclusion, we demonstrated important differences between protein and RNA expression in NSPCs and uncovered significant discrepancies in gene expression between cultured versus acutely isolated cells. Combined, our data suggest that no one method or level of analysis is sufficient for characterizing the NSPC secretome and we propose that the various proteins identified here as highly expressed in different platforms represent a putative NSPC secretome that requires subsequent validation. As stem cell based therapies are currently being developed with both cultured and endogenous cells, we recommend a multifaceted approach that includes unbiased global proteome analysis in conjunction with transcriptome analysis for the comprehensive characterization of in vitro and in vivo NSCs.

## 4. Experimental Procedures

### 4.1. Animals

Male and female C57/Bl6 mice (Jackson) aged 8-10 weeks were group housed (up to 5 per cage) in standard ventilated cages with food and water ad libitum on a 12 hour light cycle. Animals were anesthetized with intraperitoneal injections of 87.5 mg/kg ketamine / 12.5 mg/kg xylazine before perfusion and harvest of brains for primary cell derivations. This study was approved by the Institutional Animal Care and Use Committee (IACUC) at the Ohio State University.

### 4.2. Cell Culture

NSPCs were isolated from adult DGs of C57/Bl6 mice as described in (Babu et al. 2011). NSPCs were cultured on poly-D-lysine (Sigma) and laminin (Invitrogen) coated plates in Neurobasal A media (Invitrogen) with 1x B27 supplement without vitamin A (GIBCO), 1x glutamax (Invitrogen) and 20 ng/ml each of EGF and FGF2 (Peprotech). All cells used were between passage six and 15. Two separate lines were used in experiments, one from 4 pooled C57/Bl6 male mice and one from 4 pooled C57/Bl6 female mice. No inherent differences in morphology or proliferation between NSPCs isolated from males and females were found. Both lines were verified to be mycoplasma-free and to produce neurons and glia upon culture in differentiation conditions (no growth factors, data not shown).

Primary microglia used to generate conditioned media for comparison with NSPC conditioned media in VEGF ELISA were derived from one week old C57/Bl6 mice of mixed sexes. Hippocampi were isolated and pooled from 5 mice. Enzymatic digestion with 2.5 U/mL papain (Sigma), 1 U/mL dispase (StemCell Technologies), and 250 U/mL DNase (StemCell Technologies) was conducted in a 37°C water bath for 15 min. Single cell suspensions were filtered through a 70um strainer and washed with PBS (Gibco) containing 0.5% BSA (Cell Signaling Technology) before incubation with CD11b coated magnetic beads (Miltenyi) for 15 min at 4°C. Magnetic bead isolation was completed per manufacturer recommendations. All centrifugation steps were completed at room temperature at a speed of 450g for five min. Isolated cells were cultured on PDL coated plates in DMEM (Gibco) supplemented with 10% horse serum (HyClone) and 1% penicillin/streptomycin (Gibco) for three weeks prior to use.

### 4.3. Cytokine antibody array

Conditioned media was generated from three separate cultures of NSPCs, one male and two female, at 80% confluence. After 48 hours of culture, media was collected and centrifuged at 1000g for 10 min to remove large debris. For sample normalization, NSPCs for each corresponding conditioned media sample were also harvested with accutase (Invitrogen) and total protein was quantified with a BCA protein assay kit (Pierce) per manufacturer recommendations. Preparation and staining of slides were completed per manufacturer recommendations with the following modifications (RayBiotech L308 cytokine antibody array). The second dialysis step both before and after labeling of samples with biotin were conducted overnight at 4°C. Incubation of slides with biotin-labeled samples was also conducted overnight at 4°C. Incubation with Cy3-conjugated streptavidin was conducted for two hours at room temperature. After completion of the protocol, slides were completely dried before they were stored at −20°C in the 30 mL centrifuge tube provided by the manufacturer until processing. Slides were scanned by RayBiotech and raw data was analyzed according to manufacturer instructions using the RayBio Analysis Tool Software.

### 4.4. VEGF ELISA

A VEGF ELISA (R&D Systems) was performed per manufacturer instructions with cell culture supernatants. Conditioned media was generated from 2-8 biological replicates of primary mouse hippocampal microglia or adult DG NSPCs for 48 hours. Large debris was removed by centrifugation for 10 min at 1000g. VEGF standards were created with the base media for the respective cell types. Microplates were read with a microplate reader (Biotek) at 450 nm wavelength.

### 4.5. RNA isolation, cDNA library construction, and sequencing

Approximately 30,000 cells from two female and two male adult NSPCs cultures were harvested with accutase and washed with PBS before RNA isolation with the Clontech Nucleospin RNA XS Plus isolation kit (Takara 740990.10) per manufacturer protocol. RNA was also isolated from whole DGs isolated from one male and one female C57/Bl6 mouse with the same kit. RNA quality and quantity was assessed with a Qubit RNA HS assay kit (Invitrogen). All cultured samples used had a RIN value of 10 while whole DG samples had RIN values over 8. Libraries were generated with the NEBNext Ultra II Directional RNA Library prep kit (New England Biolabs) for the whole DGs and the Clontech SMART-Seq HT (Takara) kit for cultured NSPCs. Purified library products were then used in HiSeq 4000 paired-end sequencing (Illumina) to a depth of 15 - 20 million 2×150 bp clusters.

### 4.6. RNAseq analysis

Individual FASTQ files were trimmed for adapter sequences and filtered for a minimum quality score of Q20 using AdapterRemoval v2.2.0. Preliminary alignment using HISAT2 v2.0.6 was performed to a composite reference of rRNA, mtDNA, and PhiX bacteriophage sequences obtained from NCBI RefSeq. Reads aligning to these references were excluded in downstream analyses. Primary alignment was performed against the mouse genome reference GRCm38p4 using HISAT2. Gene expression values for genes described by the GENCODE Gene Transfer Format (GTF) release M14 (mouse) were quantified using the featureCounts tool of the Subread package v1.5.1 58 in stranded mode.

### 4.7. Statistics

Fluorescence intensity for each protein in the protein array was normalized to the average fluorescence intensity within each slide. Z-scores were calculated in Microsoft Excel and a heatmap of top 50 secreted factors was generated with Prism Graphpad software. For Spearman correlation with RNAseq data, fluorescence values normalized to the average fluorescence intensity within each slide were calculated in Microsoft Excel before statistical analysis and graphical representation with Prism Graphpad software.

For the VEGF ELISA, absolute protein concentrations for each sample were determined based on the standard curve and graphically represented using Prism Graphpad software.

For RNAseq analysis, published data was retrieved in Excel format from (Shin et al. 2015) and (Hochgerner et al. 2018). scRNAseq counts for each gene were averaged to generate a representative count for each population. Shin NSCs were averaged from pseudotime point 0 to 0.479. Shin nIPCs were averaged from pseudotime point 0.506 to 1. Spearman correlations were completed between RNAseq counts for the cultured NSPCs and the Shin or Hochgerner scRNAseq counts averaged over cell type for each population. For cross comparison between all datasets, counts over total counts per cell or sample were calculated and normalized to *Actb*, multiplied by a factor of 1000, before taking the log_10_ to enable graphical representation of all datasets together in a heatmap. All calculations were performed in Microsoft Excel. Spearman correlation statistics (α = 0.05) were completed for normalized scRNAseq and RNAseq datasets with Prism Graphpad software.

### 4.8. Data Availability

Processed reads for NSPC RNAseq data are provided as supplemental table 2. Upon publication, raw data files will be uploaded and publicly available in GEO.

## Acknowledgements

This work was supported by seed funding from the Chronic Brain Injury and Discovery Themes at The Ohio State University to EDK and R00NS089938 to EDK from the NIH/NINDS.

